# Myocardial cGMP-PKG1α dysregulation contributes to VT pathogenesis in type II diabetes and metabolic syndrome

**DOI:** 10.1101/2025.02.12.638003

**Authors:** Xuehong Cao, Yali Zhang, Audrey Tripp, Rachel Steinhauer, Pei-wen Liu, Mark J. Aronovitz, Greg L. Martin, Bo Wang, Abdullah Alissa, Mossab Aljuaid, Justin Ho, Tina Phan, Christopher Madias, Robert M. Blanton, Jonas B. Galper

## Abstract

**Background:** Type-II diabetes (DMII) and metabolic syndrome increase ventricular arrhythmia and sudden cardiac death risk.

**Objectives:** To identify signaling mechanisms through which DMII and metabolic syndrome promote ventricular tachycardia (VT).

**Methods:** We performed ventricular programmed stimulation on leptin receptor mutant (Db/Db) mice with DMII, high fat high sucrose (HFHS)-fed mice with metabolic syndrome, and cGMP-dependent Protein Kinase 1α (PKG1α) leucine zipper mutant (LZM) mice, which do not have DMII or metabolic syndrome but have disrupted PKG1α signaling.

**Results:** During ventricular programmed stimulation, Db/Db and HFHS-fed mice displayed increased VT and T-wave alternans. Cardiomyocytes from these mice displayed early afterdepolarizations. Both models demonstrated decreased heart rate response to parasympathetic inhibition, indicating autonomic dysfunction. cGMP, which mediates cardiac parasympathetic stimulation, was reduced in LVs of Db/Db and HFHS-fed mice. Conversely, cGMP augmentation with soluble guanylate cyclase stimulation (riociguat) or phosphodiesterase 5 inhibition (sildenafil) reduced VT inducibility. PKG1α LZM mice had normal autonomic responsiveness, but excess VT inducibility. Db/Db, HFHS, and LZM mice each demonstrated hyperactivated myocardial glycogen synthase kinase3β (GSK3β). Further, GSK3β inhibition with TWS119 abolished inducible VT in these mice. Diastolic cytosolic Ca^2+^ reuptake slope decreased in cardiomyocytes from all models, while GSK3β inhibition with TWS119 reversed this effect. Phospholamban (PLB), which inhibits sarcoplasmic/endoplasmic reticulum Ca^2+^ ATPase 2a-mediated Ca^2+^ reuptake, was hyperactivated/hypophosphorylated in HFHS-fed and LZM mice, and this was reversed by TWS119.

**Conclusions:** These findings identify cGMP reduction as driving GSK3β hyperstimulation, calcium dyshomeostasis, and VT in DMII and metabolic syndrome. Pharmacological modulation of these pathways opposes VT pathogenesis.

## Introduction

Sudden cardiac death (SCD) remains a leading cause of cardiovascular death^1^. SCD typically arises from lethal ventricular arrhythmias including ventricular tachycardia (VT). VT and SCD occur either in the setting of acute myocardial infarction (MI), or in patients with chronic heart failure with reduced LV ejection fraction (HFrEF). Improvements in acute revascularization and in inpatient telemetric monitoring have greatly reduced SCD in the immediate post-MI period^2^. Similarly, placement of implantable cardiac defibrillators in patients with HFrEF and LV ejection fraction ≤ 35% has improved survival by reducing VT death^3^. Despite these advances, VT remains a significant cause of death in patients with mildly reduced or even normal LVEF^1^. Thus, identifying and understanding additional mechanisms of the pathogenesis of VT has the potential to improve prevention of SCD in patients.

Diabetes (DM) and metabolic syndrome represent poorly understood risk factors for SCD and VT. Both type 1 and 2 DM (DMI, DMII) as well as insulin resistance and metabolic syndrome, increase SCD risk in patients^4–6^. Epidemiological studies support that DMII independently increases SCD risk both in patients with known heart disease, but also in patients without known underlying heart disease^7,8^. In a recent study of patients with DMII who died under 50 years of age, SCD accounted for the most frequent cause of death. Approximately half of DMII patients with SCD had no evidence of CAD on autopsy^7^. Thus, available clinical evidence supports that in addition to effects on CAD and other cardiovascular risk factors, DM and metabolic syndrome serve as an independent risks for SCD.

Experimental models further support the conclusion that DM acts as a VT substrate. For example, our previous work demonstrated that the Akita mouse model of type 1 diabetes displays increased susceptibility to VT in response to programmed electrical stimulation even in the basal state, in the absence of acute MI^9^. Several experimental studies also indicate increased VT susceptibility in animal models of DMII^10,11^. These combined population and experimental studies support an independent effect of DM on susceptibility to VT. Impaired glucose tolerance and metabolic syndrome, like DMII, increase SCD risk in patients^4–6^, suggesting that shared mechanisms exist for VT predisposition in DM and insulin resistance. However, the molecular abnormalities through which DMII promotes VT susceptibility independent of associated cardiovascular risk factors remain unknown.

Autonomic dysfunction is a shared complication of DMI, DMII, and insulin resistance^12^ and has been identified as contributing to arrhythmogenesis. In patients with DMII, autonomic dysfunction further increases SCD risk ^4^. Experimental studies demonstrate that parasympathetic dysfunction contributes to inducible and spontaneous post-MI VT in DM1 due in part to excess unopposed sympathetic input to the heart.^9,13,14^. In the leptin receptor mutant (Db/Db) model of DMII, myocardial autonomic dysregulation promotes VT as assessed by an *ex vivo* inducible arrhythmia model^10^. In the myocardium, parasympathetic signaling is normally mediated by the elevation of the second messenger cyclic guanosine monophosphate (cGMP), consistent with the finding that parasympathetic dysfunction is associated with reduced myocardial cGMP^15^.

However, a causal role of reduced cGMP signaling in the susceptibility to VT remains unknown. Moreover, the specific role of derangements in cGMP signaling in the pathogenesis of VT in DMII and metabolic syndrome has not been studied.

In the present study we therefore examined the dysregulation of myocardial cGMP in mouse models of DMII and insulin resistance and investigated the role of the cGMP-dependent protein kinase 1 (PKG1) in modulating the predisposition to the development of VT, using a protocol for programed ventricular stimulation. We specifically studied the Db/Db mouse harboring leptin receptor mutations which produce obesity, DMII, and metabolic heart disease^16^, and the high-fat high-sucrose (HFHS)-fed mouse which by contrast, does not have overt DM, but develops obesity, metabolic syndrome, and insulin resistance^17^. We measured myocardial cGMP content and tested the anti-arrhythmic effects of pharmacological augmentation of cGMP with the cGMP-specific phosphodiesterase 5 (PDE5) inhibitor sildenafil and the guanylate cyclase stimulator ricociguat on VT in the Db/Db and HFHS models. To study the direct role of PKG1, we used the PKG1α leucine zipper mutant (LZM) mouse, which has normal cGMP levels, does not have diabetes or metabolic syndrome, but lacks the normal PKG1α response to cGMP due to point mutations which disrupt kinase substrate interactions^18^. This allows the specific investigation of cGMP-PKG1 signaling without being confounded by other effects of DM.

To establish the molecular mechanism for increased inducibility of VT in the above models, we studied Ca^2+^ transients in cardiomyocytes and dysregulation of Ca^2+^ handling proteins in ventricular extracts from these mice. We also investigated QRS/T-wave alternans and Ca^2+^ alternans in the above models, as these phenomena occur due to Ca^2+^ dyshomeostasis and both correlate with and contribute to lethal ventricular arrhythmia in humans^19–25^. Finally, we determined contribution of glycogen synthase kinase 3β (GSK3β) signaling in mediating the cGMP-PKG1 effect on VT phenotype.

## Methods

### Institutional approval

All mouse studies were approved in accordance with the Institutional Animal Care and Use Committee at Tufts University, Protocol B2022-159.

### Mouse models

The Db/Db leptin receptor mutant mice were obtained from Jackson Laboratories (strain 000697). Male mice were studied at age 4-6 months. We used Db/+ age-matched littermates which lack the Db/Db phenotype as controls. The high fat high sucrose (HFHS)-fed mice were treated for at least 24 weeks, starting at 8 weeks of age on 58kcal% fat, 656 kcal sucrose, and 5% insoluble fiber, cellulose (Research Diets, D23071405). Male wild type littermates received a control diet containing 4.8% insoluble fiber, 10kcal% fat and 0 sucrose (Research Diets D23071406). The PKG1α leucine zipper mutant (LZM) mice have been described previously^18^. We compared male homozygous LZM mice with wild type littermate age-matched controls at 4-6 months of age.

### Programmed Stimulation

We performed the intracardiac programmed stimulation protocol as described previously^9^. Briefly, mice underwent light anesthesia with 2% isoflurane. A surface frontal plane ECG was obtained using 29-gauge electrodes placed subcutaneously in each limb. A 1.1-Fr electrophysiology octopolar catheter (EPR-800, Millar Instruments, Houston, TX) was inserted via the right jugular vein and advanced to the right ventricle. Next, the position and capture were confirmed through intracardiac and surface ECG tracings. Following pacing at 90ms hearts were subjected to premature stimuli at decreasing S1-S2 intervals until refractory, followed by a second S2 and third S3 premature stimulus until refractory. Finally, mice were subjected to burst pacing at rates of 40, 50, 60, 70, 80 and 90ms. Total duration of induced ventricular tachycardia in response to these stimuli was reported^9^. The threshold for development of alternans was determined as the slowest pacing rate at which T-wave alternans appeared.

### Heart rate response to atropine

Mice received surgically implanted ECG telemeters as described^47^, followed by recovery for at least 3 days. Mice were treated with intraperitoneal injections of atropine, 0.5 mg/kg and the change in mean heart rate at 4 minutes post-injection was compared with the heart rate averaged over the minute prior to injection. LZM mice were compared with the same control mice as the Db/Db and HFHS mice.

*cGMP concentration* in LV tissue was immediately snap frozen in liquid nitrogen following sacrifice. Approximately 30mg of tissue was then crushed into a fine powder over dry ice using a pre-chilled cryo tissue grinder. On ice, 6% trichloroacetic acid was added to samples (10µL/mg of tissue) followed by vortexing for 10 seconds. Proteins were precipitated out by centrifuging at 2000 x G for 15 minutes at 4°C and the supernatant was transferred to a 15 mL falcon tube on ice. Trichloroacetic acid in the supernatant was extracted by addition of approximately five volumes of diethyl ether. Samples were mixed by gentle rocking, then allowed to equilibrate into two layers. The top layer was discarded, and the sample washed a further three times by repeat additions of diethyl ether. After the final wash, the bottom layer was transferred to a microcentrifuge tube, snap frozen in liquid nitrogen and later lyophilized. Samples were resuspended in assay buffer and cGMP concentration determined in duplicate using the cGMP Enzyme Immunoassay Biotrak (EIA) system (RPN226, GE Healthcare, Amersham, UK), according to protocol #2.

### Drug administration

Riociguat (AmBeed, A255332) was administered by oral gavage at 10 mg/kg once daily for 7 days. Sildenafil (Tocris 3784 or Sigma-Aldrich SML3033) was administered by intraperitoneal injection at 10 mg/kg once daily for 7 days. TWS199 (MEMD Millipore Corp, USA, 361554) was administered at 20 mg/kg by intraperitoneal injection once daily for 7 days. The response of each agent was compared with saline vehicle.

### Immunoblotting

LV tissue lysates were prepared as described^14^. We used the following antibodies as described in **Supplemental Table 1**.

### CM isolation and analysis

Adult mouse ventricular cardiac myocytes were isolated from mice as described previously^47^, plated on laminin-coated cover slips followed by equilibration in physiological calcium, and loaded with FURA-2-AM (1µM, Invitrogen) for 30 minutes room temperature. Cells were placed in a chamber and paced via field stimulation at 1Hz, 2Hz, 3Hz, 4Hz and 5Hz. Sarcomere shortening and Ca^2+^ transients were determined using a dual excitation spectrometer and video edge detection (Ionoptix) as described previously^47^). Data were plotted using IonWizard. Ca^2+^ reuptake rate was determined from the slope of the recovery curve, Tau. Ca^2+^ alternans threshold was determined as the lowest pacing rate at which cells develop Ca^2+^ alternans. We defined alternans as a 20% variation in beat-to-beat amplitude.

### Early afterdepolarization (EAD) recording in cardiac myocytes

EADs were recorded in cardiomyocytes using whole-cell patch clamp in the mode of current clamp at room temperature. Recordings were performed with an Axon 200B amplifier and DigiData 1440 (Molecular Devices) and analyzed with Clampfit 11.2. The bath solution consisted of (in m<u>M</u>): 137 NcAl, 5.4 KCl, 1 CaCl_2_, 0.5 MgCl_2_, 0.33 NaH_2_PO_4_, 5 HEPES, and 5.5 glucose, with a pH of 7.4 and an osmolality of 300 mOsm. Cardiomyocytes were paced by brief (2 ms) depolarizing pulses at frequencies of 0.1, 0.2, 1, 2, 3, and 5 Hz to induce and study EADs.

### Statistical analysis

Average values are given as the mean ±SEM. Statistical differences between mean values were calculated by Student’s T test, Welch T test, or Mann-Whitney test as described in the figure legends. For multiple group comparisons, 1-way ANOVA was used, followed by multiple comparison testing.

## Results

### Db/Db and HFHS models of DMII and metabolic syndrome display increased VT in response to programmed electrical stimulation and increased incidence of T wave alternans

Measures obtained from surface ECG recording in HFHS-fed and Db/Db mice are displayed in **Table 1**. HFHS-fed mice displayed mildly increased P wave duration compared to control-diet treated, and had a reduced R-R interval, indicating faster heart rate. We examined VT inducibility in Db/Db and HFHS mice using ventricular programmed electrical stimulation. Programmed electrical stimulation of the ventricle predicts both SCD risk and VT events in patients following myocardial infarction^26,27^. We previously optimized a unique technique for programmed ventricular stimulation in mice with DMI utilizing a catheter with electrodes spatially designed to stimulate and record from the mouse heart to investigate and identify molecular mechanisms predisposing to VT^9,28^. The total duration of VT in response to single S1, double S2, and triple S3 stimulation was 3.2 ± 0.9 sec in Db/Db, (n= 23) vs. 0.3 ± 0.2 sec in Db/+, (n = 14, p<0.05).

**Table 1.** Surface ECG intervals in experimental groups.

| **Condition** | **N** | **P (ms)** | **PR (ms)** | **RR (ms)** | **HR (min<sup>-1</sup>)** |
| --- | --- | --- | --- | --- | --- |
| Control Diet | 7 | 11.62 ± 1.06 | 54.33 ± 4.96 | 146.57 ± 8.18 | 416.37 ± 21.10 |
| HFHS Diet | 16 | 14.56 ± 0.42 | 54.63 ± 1.21 | 137.08 ± 3.95 | 442.90 ± 12.11 |
| p |  | 0.005 | ns | ns | ns |
| HFHS Vehicle (for Sildenafil) | 5 | 12.00 ± 0.42 | 46.87 ± 2.67 | 132.00 ± 8.42 | 462.42 ± 31.01 |
| HFHS Sildenafil | 6 | 10.94 ± 0.51 | 48.11 ± 1.74 | 134.67 ± 8.57 | 455.02 ± 29.86 |
| P |  | ns | ns | ns | ns |
| HFHS Vehicle (for Riociguat) | 5 | 13.93 ± 0.31 | 52.73 ± 1.96 | 136.60 ± 16.92 | 461.01 ± 44.44 |
| HFHS Riociguat | 5 | 11.60 ± 0.96 | 54.60 ± 0.96 | 126.20 ± 3.53 | 476.82 ± 12.42 |
| P |  | 0.050 | ns | ns | ns |
| Db/+ | 4 | 11.08 ± 10.3 | 45.75 ± 1.15 | 172.33 ± 21.34 | 364.12 ± 43.66 |
| Db/Db | 19 | 13.35 ± 0.62 | 48.05 ± 2.00 | 138.23 ± 5.78 | 447.58 ± 18.41 |
| P |  | ns | ns | 0.040 | ns |
| Db/Db Vehicle (for Sildenafil) | 4 | 14.75 ± 1.01 | 51.00 ± 3.88 | 138.25 ± 6.33 | 436.72 ± 19.85 |
| Db/Db Sildenafil | 5 | 14.13 ± 0.77 | 54.27 ± 2.80 | 147.73 ± 7.59 | 410.55 ± 21.52 |
| P |  | ns | ns | ns | ns |
| Db/Db Vehicle (for Riociguat) | 14 | 13.58 ± 0.83 | 55.61 ± 5.38 | 183.22 ± 12.86 | 337.74 ± 19.07 |
| Db/Db Riociguat | 11 | 11.88 ± 0.77 | 48.97 ± 1.97 | 145.85 ± 6.02 | 418.19 ± 16.58 |
| P |  | ns | ns | 0.024 | 0.005 |

The total duration of VT was 14.6 ± 3.0 sec in HFHS fed mice (n=14), vs. 2.6 ± 2.1 sec in mice on control diet (n=7, p<0.05) (**Figure 1A-1C**). Burst pacing had no effect on the development of VT in either experimental group. Female mice on HFHS diet displayed no inducible VT, resulting in a statistically significant difference in VT duration compared with male HFHS mice. For this reason, subsequent studies investigated males only.

**Figure 1.**
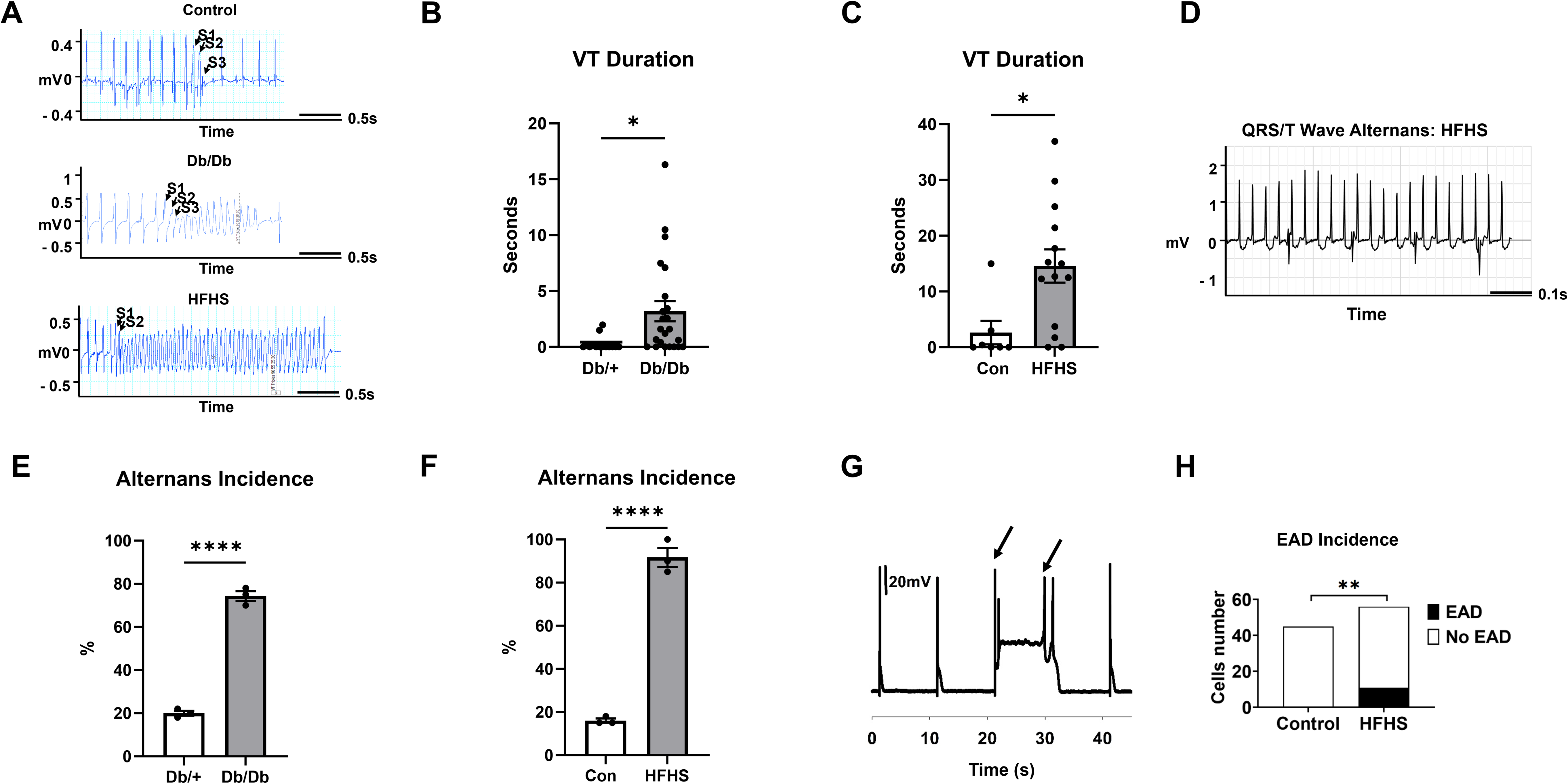
Increased ventricular tachycardia inducibility and T-wave alternans in mice with type II diabetes and metabolic syndrome. (A) Representative intracardiac tracings demonstrating VT in Db/Db and HFHS, but not control mice, during programmed stimulation. (B) Total VT duration in response to programmed stimulation and burst pacing in Db/Db, Db/+ controls and (C) HFHS versus control diet. *, p<0.05 by Student’s unpaired T Test. (D) Representative T-wave alternans in response to burst pacing in HFHS fed mouse. (E) T-wave alternans incidence in Db/+, Db/Db, and (F) wild type mice on control and HFHS diet, during 20 Hz interval burst pacing. (G) Early afterdepolarizations (EADs) in action potential from ventricular myocyte of HFHS-fed mouse paced at 100 Hz. Arrows denote EADs. (H) EAD incidence in control and HFHS diet-fed mouse ventricular myocytes. **, p<0.01 by Chi-squared test, n= 0/45 cells with EADs in 5 control; 11/56 cells in 7 HFHS-fed mice.

Further analysis of the surface ECG in these mice during burst pacing demonstrated a rate-related beat-to-beat alteration voltage consistent with QRS/T-wave alternans (**Figure 1D**). Compared with respective Db/+ or control diet controls, the Db/Db and HFHS mice displayed increased incidence of QRS/T-wave alternans with pacing rates of 70, 60, and 50ms (**Figure 1E, 1F**). At 50ms pacing rates both Db/Db and HFHS mice displayed statistically significantly increased incidence of T wave alternans compared with controls (74.3% in Db/Db mice vs 20% in Db/+ mice, p<0.001, and 91.7% in HFHS fed mice compared with 16% in mice fed a control diet, p<0.0001).

These data support a temporal or spatial dispersion of myocyte repolarization and/or abnormalities of Ca^2+^ cycling which might play a role in the predisposition to VT in these mice. To examine this further, we determined whether cardiomyocytes from these mice develop early afterdepolarizations (EADs) in response to increasing pacing rates. We measured action potentials in adult ventricular myocytes isolated from HFHS-fed mice at pacing rates of 0.1 to 5Hz. Data summarized in **Figure 1H** demonstrate typical early afterdepolarizations at 0.1 Hz characterized by reversal of the action potential before repolarization is complete and a second EAD preceding the subsequent paced beat. Although 0 of 45 CMs from 5 mice on the control diet developed EADs, 7 of 56 cells from 7 mice on a HFHS diet (18%) developed EADs, (p<0.01 by chi-squared test, **Figure 1G, 1H**). EADs, which cause ectopic beats and lethal arrhythmias reflect decreased repolarization reserve and/or an associated increase in cytosolic calcium ^29^. These findings support the conclusion that in mice with DMII or metabolic syndrome, the LV develops increased susceptibility to VT which may be associated with Ca^2+^ dyshomeostasis.

### Blunted parasympathetic response and decreased cGMP levels in hearts of Db/Db and HFHS fed mice

Diabetic autonomic neuropathy, characterized initially by parasympathetic dysfunction, is a major complication of both type I and type II diabetes^9,13^. We therefore determined whether parasympathetic signaling was blunted in Db/Db and HFHS fed mice. We determined the heart rate response to a 0.5 mg/kg injection of the acetylcholine receptor antagonist atropine in Db/Db and in HFHS mice, as a readout for parasympathetic function. Compared with wild type control mice, the HFHS and Db/Db mice each displayed reduction in atropine-induced heart rate increase (**Figure 2**) (relative beat-per-minute increase: 51 ± 5% in mice fed a control diet, 20 ± 3% in HFHS fed mice and 23 ± 10% in Db/Db mice, p<0.01), indicating a markedly reduced parasympathetic heart rate response in the latter 2 models.

**Figure 2.**
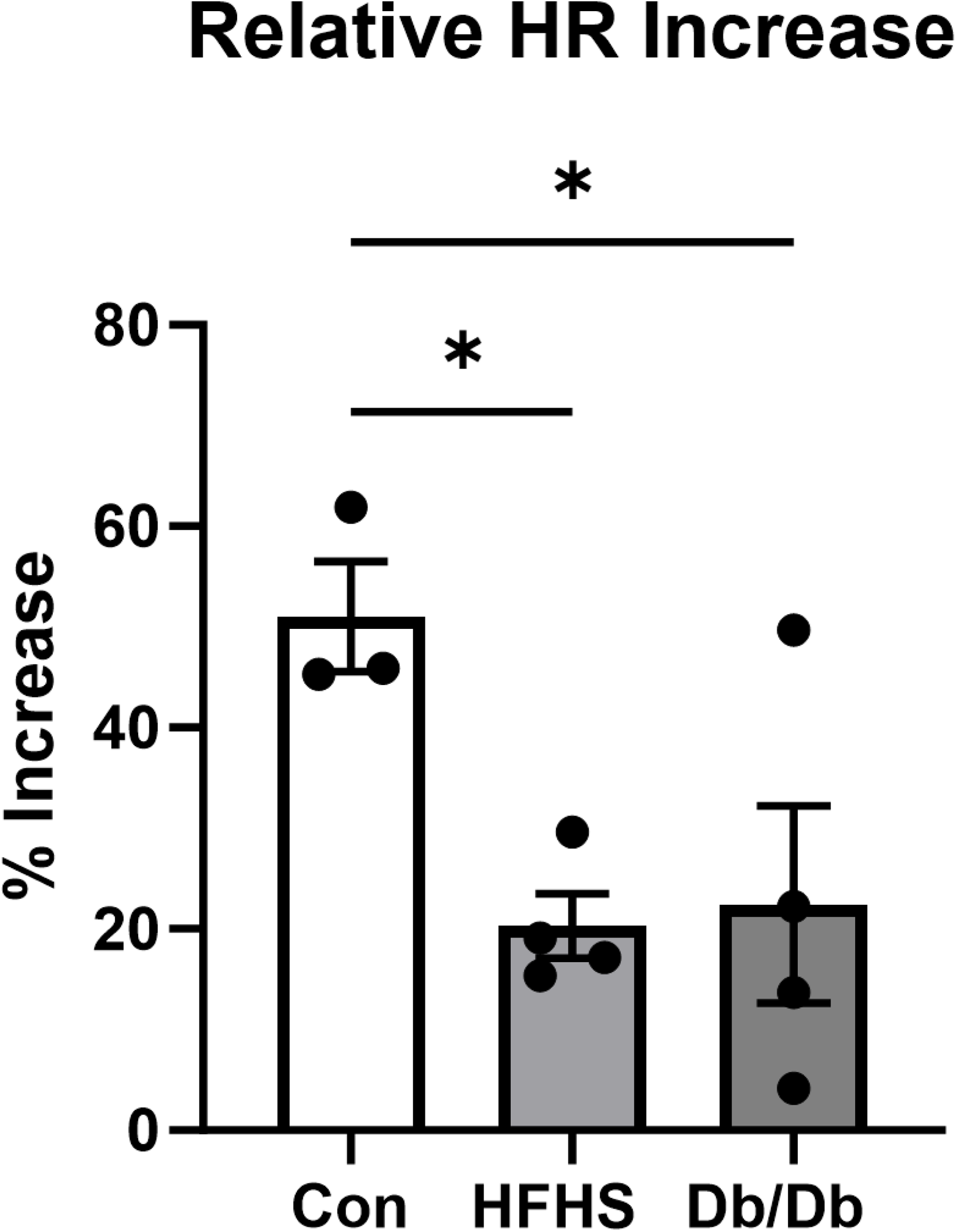
Parasympathetic dysfunction in mice with type 2 diabetes or metabolic syndrome. Relative increase in heart rate (HR) measured by implantable EKG transmitters in unanesthetized mice following intraperitoneal injection of 0.5 mg/kg atropine in mice fed control (Con) diet, high fat high sucrose (HFHS) diet, or Db/Db mice. *, p<0.05 by 1-way ANOVA with Dunnett’s multiple comparisons test.

Parasympathetic stimulation promotes myocardial cGMP synthesis via the stimulation of the M2 muscarinic receptor^15^. Given the observed decrease in cardiac parasympathetic function in the Db/Db and HFHS mice, we tested for concomitant reduction in myocardial cGMP. cGMP concentration was reduced in LV extracts of both Db/Db and HFHS-fed mice, compared to control (**Figure 3A, 3C**). cGMP decreased 21 ± 4% in Db/Db vs Db/+ (p<0.01), and 29 ± 8% in HFHS-fed mice vs mice fed a control diet, (p<0.05). Additionally, LV protein expression of the cGMP-specific phosphodiesterase 5 (normalized to β-actin) was increased versus control both in Db/Db and HFHS (1.6 ± 0.04-fold in Db/Db vs. Db/+ mice, p<0.0001; 1.7 ± 0.2-fold increase in HFHS fed mice vs. control diet, p<0.05) (**Figure 3B, 3D**). Thus, cGMP levels in Db/Db and HFHS fed mice were decreased both at the level of synthesis via decreased muscarinic stimulation and increased turnover via increased PDE5-associated degradation.

**Figure 3.**
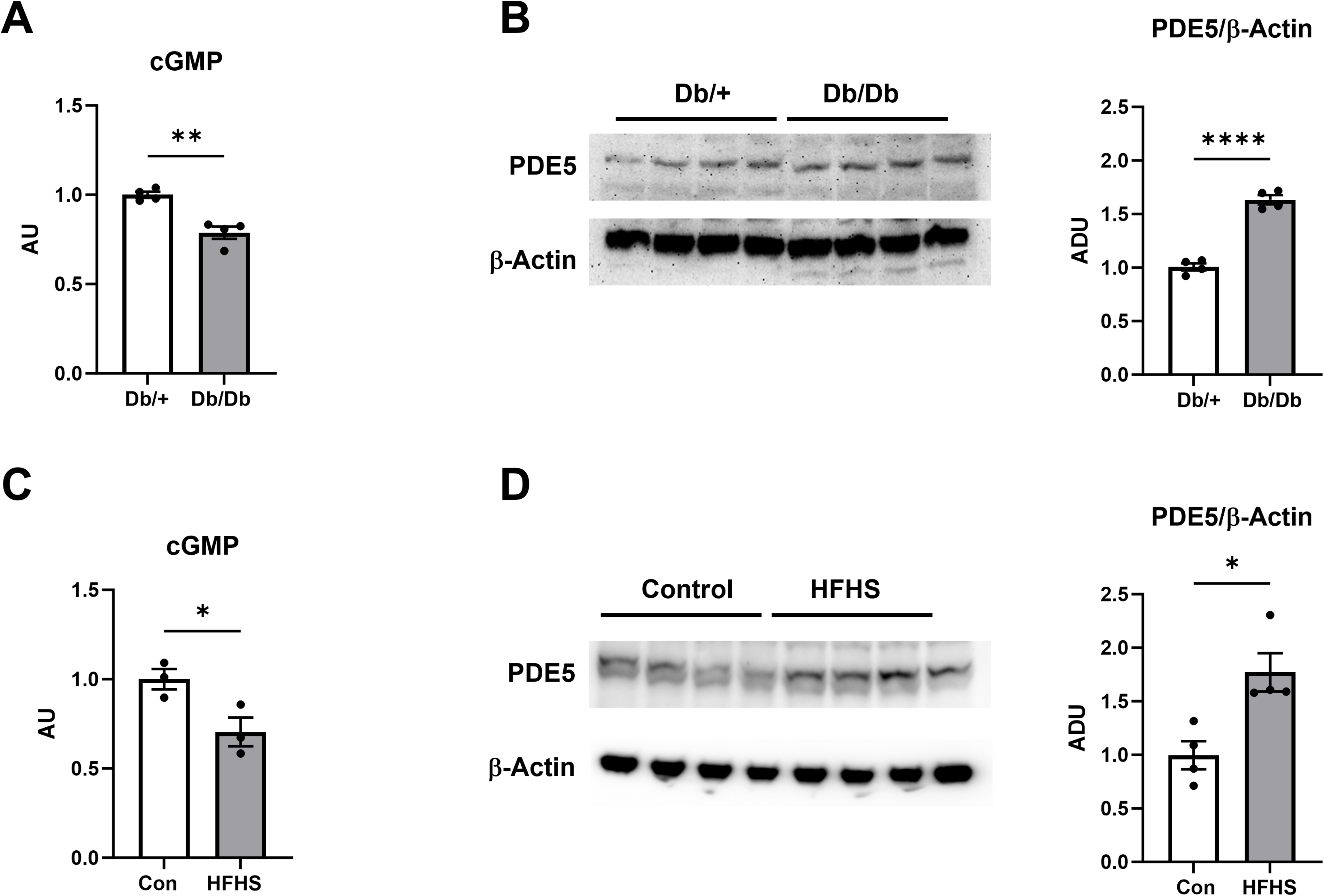
Reduced cGMP and increased cGMP-catabolizing phosphodiesterase 5 in LV tissue of mice with type II diabetes mellitus and metabolic syndrome. (A), cGMP levels and (B), representative immunoblot and densitometric analysis of phosphodiesterase 5 (PDE5) normalized to β-Actin in LV tissue from Db/+ and Db/Db mice. (C), cGMP levels and (D), representative immunoblot and densitometric analysis of phosphodiesterase 5 (PDE5) normalized to β-Actin in LV tissue from mice fed a high fat high sucrose (HFHS) diet or control diet (Con). *, p<0.05; **, p<0.01; ****, p<0.0001 by Student’s unpaired T test.

### Stimulation of cGMP synthesis with riociguat or inhibition of cGMP degradation with the PDE5 inhibitor sildenafil reverses inducible VT in Db/Db and HFHS fed mice

To determine whether the reduction in cGMP signaling in the LV of Db/Db and HFHS fed mice played a role in their predisposition to VT, we treated Db/Db and HFHS-fed mice with the cGMP-augmenting soluble guanylate cyclase stimulator riociguat (10 mg/kg for 7 days), followed by programmed ventricular stimulation as above. Riociguat and sildenafil effects on ECG intervals are shown in **Table 1**. Riociguat treatment reduced duration of VT in both the Db/Db mice (**Figure 4A**) and in the HFHS-fed mice (**Figure 4B**) compared with respective vehicle-treated controls. Specifically, total duration of VT in Db/Db mice decreased from: 4.1 ± 1.5 sec in vehicle-treated (n=13) to 0.9 ± 0.5 sec in riociguat-treated (n=15) mice (p<0.05).

**Figure 4.**
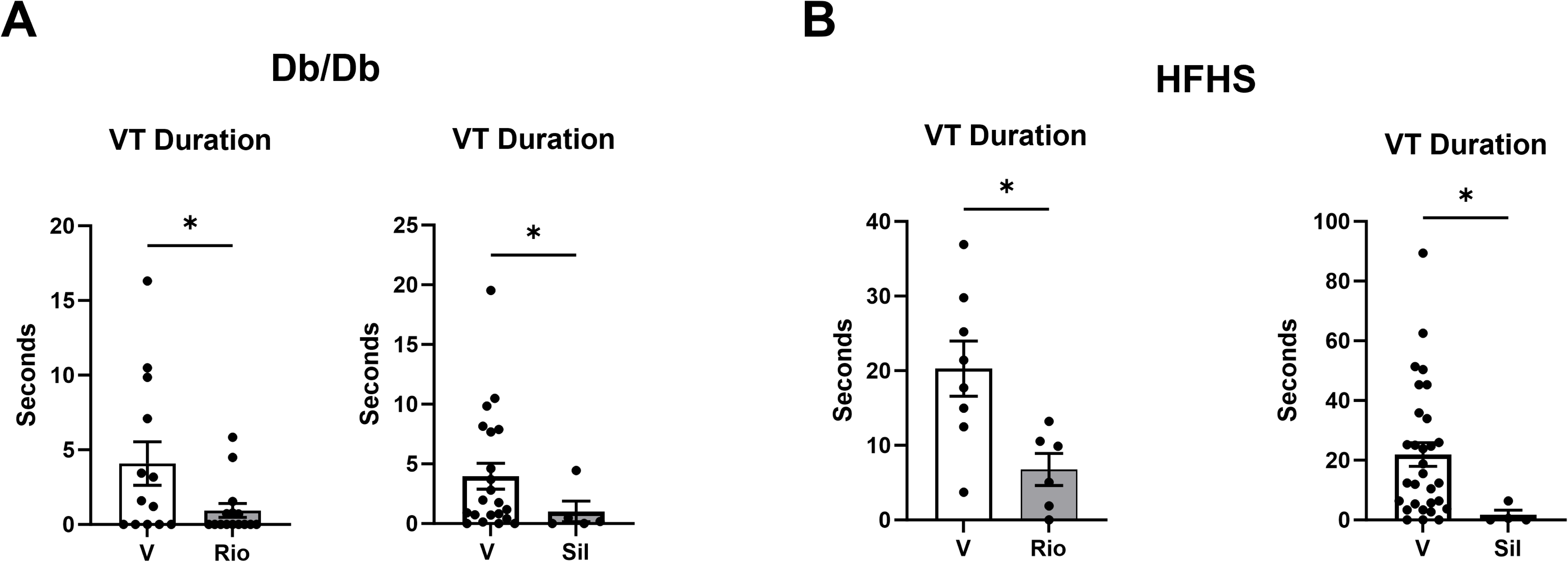
cGMP augmentation with the soluble guanylate cyclase stimulator riociguat or the phosphodiesterase 5 inhibitor sildenafil reduces duration of ventricular tachycardia in mice with type II diabetes and metabolic syndrome. Development of ventricular tachycardia (VT) in response to programmed ventricular stimulation in (A) Db/Db mice and (B) high fat high sucrose-fed mice pre-treated for 7 days with vehicle (V), riociguat (Rio) 10 mg/kg, or sildenafil 10 (Sil) 10 mg/kg. *, p<0.05 by Student’s unpaired T test (riociguat graphs) or in sildenafil-treated graphs, by Welch’s T test (Db/Db) or Mann-Whitney test (HFHS).

Duration of VT in HFHS fed mice decreased from 20.3 ± 3.7 sec in vehicle-treated (n=8) to 6.8 ± 2.2 sec in riociguat-treated (n=6) mice, (p<0.05). We tested a second cGMP-augmenting drug, the phosphodiesterase 5 inhibitor sildenafil (10 mg/kg for 7 days), and again observed in HFHS-fed mice that sildenafil treatment reduced cumulative VT duration, compared with vehicle-treatment. Total duration of VT decreased from 4.0 ± 1.1 sec in vehicle-treated (n=21) Db/Db to 1.0 ± 0.9 sec in sildenafil-treated (n=5) Db/Db mice (p<0.05). Similarly, in HFHS-fed mice duration of VT decreased from 21.9 ± 4.0 sec in vehicle-treated (n=30) to 1.7 ± 1.6 sec in sildenafil-treated (n=4) mice (p<0.05).These findings indicate that augmenting cGMP levels opposes the predisposition to VT in the Db/Db and HFHS fed mice.

### Disruption of PKG1α activity in the absence of diabetes and metabolic syndrome promotes inducible VT and T wave alternans in vivo

We next directly tested the role of the cGMP-dependent protein kinase 1α (PKG1α) in modulating VT *in vivo*. PKG1 is the primary cGMP effector in the LV, and cGMP binding to PKG1α stimulates PKG1α kinase function^30^. To examine the role of PKG1α signaling in opposing the predisposition to VT, we performed programmed ventricular stimulation on PKG1α leucine zipper mutant (LZM) mice. These mice harbor a series of point mutations within the PKG1α leucine zipper protein-protein binding domain, producing an enzyme which retains kinase function but cannot bind to LZ-dependent substrates. Importantly, the LZM mice have a normal metabolic phenotype and do not display evidence of diabetes or insulin resistance^31^. The LZM mouse has normal levels of cGMP and does not yet demonstrate ventricular hypertrophy or LV dysfunction at 12-14 weeks of age^18,32,33^. Unlike the Db/Db and HFHS-fed mice, LZM mice did not manifest autonomic dysfunction. Specifically, atropine treatment increased heart rate normally in LZM mice from 535 ± 42 beats/minute to 754 ± 15 beats/minute post-injection (n=5, p<0.01) (**Figure 5A**). Similarly, while Db/Db and HFHS-fed mice had blunted heart rate response to atropine compared with WT control mice (**Figure 2**), the LZM mice had a normal increase in heart rate when compared to the same WT mice (51.0 ± 5.4% in WT controls vs. 44.2 ± 10.4% in LZM mice, p=0.65). In contrast to the normal heart rate response to atropine, LZM mice developed increased inducible VT in response to programmed ventricular stimulation, with a total duration of 18.8 ± 7.0 seconds in LZM (n=6) vs. 0 ± 0 seconds in WT (n=4) mice (p<0.05, **Figure 5B**). Furthermore, like Db/Db and HFHS fed mice, LZM mice developed T-Wave alternans in response to pacing intervals at 60ms (**Figure 5C**). Specifically, 0 of 4 WT littermates compared with 5 of 6 LZM mice developed alternans (**Figure 5D**, p<0.001 by chi-square analysis).

**Figure 5.**
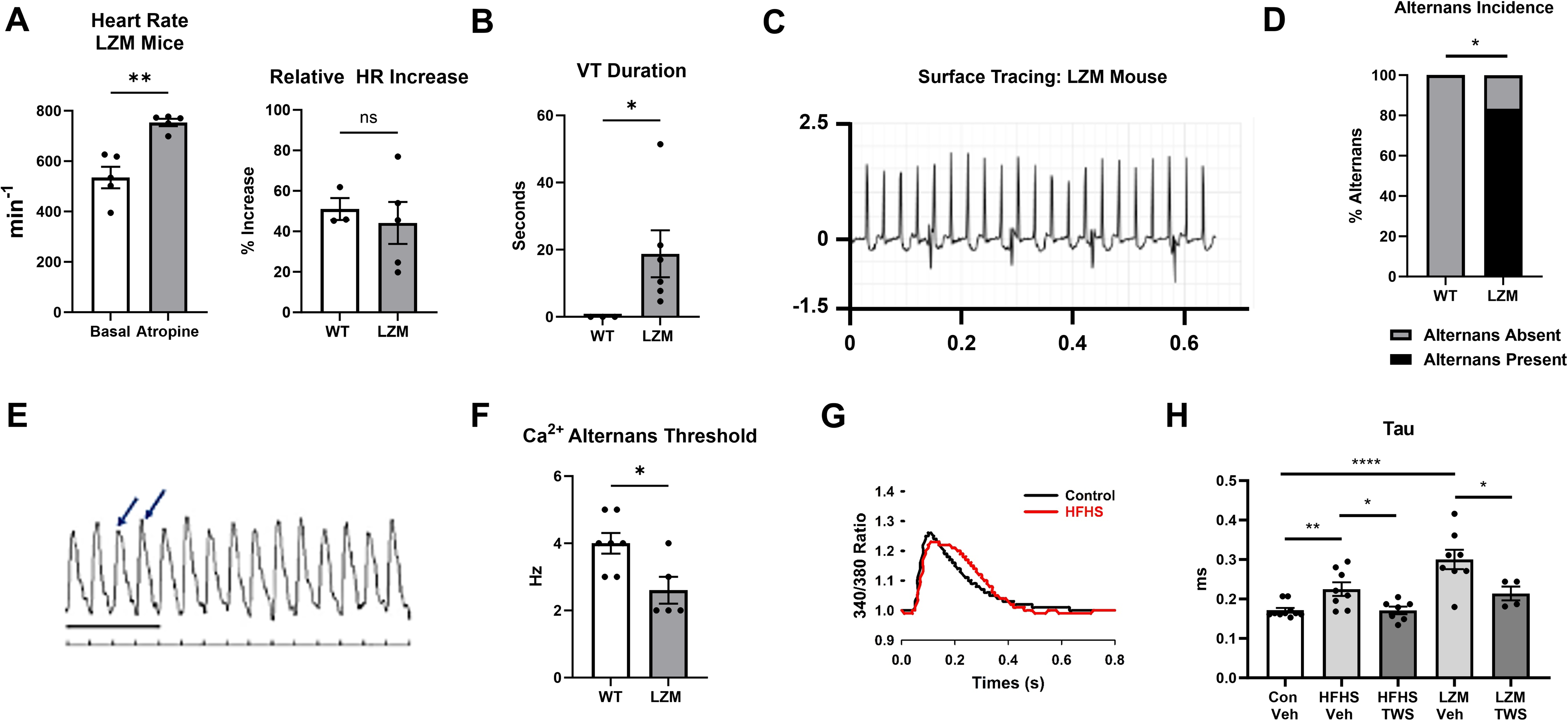
cGMP-dependent protein kinase 1α mutation promotes ventricular tachycardia and calcium dyshomeostasis independently of diabetes or metabolic syndrome. (A) Heart rate at baseline or 4 minutes after 0.5 mg/kg atropine injection in PKG1α leucine zipper mutant (LZM) mice, and atropine-induced percentage heart rate increase versus wild type. (B) VT duration during programmed stimulation in LZM and WT littermates. (C) Representative surface recording and (D) T-wave alternans incidence at 16.7 Hz pacing in LZM or WT mice. Alternans present in 0/4 WT, 5/6 LZM. *, p<0.01 by chi-squared test. (E) Ca^2+^ transients from LZM ventricular myocytes demonstrating alternans at 3 Hz. (F) Ca^2+^ alternans threshold in ventricular myocytes from LZM and WT mice across increasing pacing rates. (G) Representative Ca^2+^ transient in ventricular myocytes from WT and LZM mice. (H) Time constants of Ca^2+^ reuptake, tau, in ventricular myocytes from: WT on control (Con) or HFHS (HFHS) diet, and LZM mice; injected daily with vehicle or 20 mg/kg of GSK3β inhibitor TWS119 for 7 days. *, p<0.05; **, p<0.01; ***, p<0.001, and ****, p<0.0001 by Student’s unpaired T test.

Given the relationship between the development of T-wave alternans and abnormalities of Ca^2+^ homeostasis^34^, the observation of T-wave alternans in HFHS-fed, Db/Db, and LZM mice suggested that a common mechanism of Ca^2+^ dyshomeostasis might account for the predisposition to VT development in these mouse models. To test this hypothesis, adult ventricular myocytes isolated from LZM, HFHS-fed and WT mice were paced at increasing rates. Cells from LZM and HFHS fed mice developed Ca^2+^ alternans **Figure 5E**. Furthermore, the average pacing threshold for the development of Ca^2+^ alternans was reduced in LZM mice (2.6 ± 0.4 Hz, n=5) compared with WT mice (4 ± 0.3 Hz, n=7, p<0.05) (**Figure 5F**).

The presence of Ca^2+^ alternans suggested that shared abnormalities of intracellular calcium SR reuptake in LZM and HFHS mice might contribute to the pathogenesis of VT^34^. We therefore examined Ca^2+^ transients and the time constant of cytosolic Ca^2+^ reuptake in ventricular myocytes loaded with the Ca^2+^ sensitive dye Fura-2. Ca^2+^ transients from HFHS CMs (**Figure 5G**), demonstrated reduced slope of the Ca^2+^ reuptake curve, supporting decreased rate of sarcoplasmic reticulum Ca^2+^ from the cytosol. Quantitative comparison of Ca^2+^ transients from ventricular myocytes from HFHS fed and LZM mice with myocytes from mice fed a control diet or WT mice respectively, further demonstrated increased time constant of cytosolic Ca^2+^ decline (tau). Specifically, tau values in cardiomyocytes from HFHS fed mice were 0.23 ± 0.02 ms, n=7 vs 0.17 ± 0.01 ms in cells from mice fed a control diet (n=10, p<0.01). In cells from LZM mice (n=8) tau was 0.30 ± 0.03 ms, (p<0.0001) vs control (**Figure 5H**). These findings indicate that metabolic syndrome and PKGα dysfunction produce similar slowing in diastolic Ca^2+^ sarcoplasmic reticulum reuptake in CMs, which would predispose to VT.

### Disinhibition of GSK3β signaling in the LVs of diabetic, metabolic syndrome, and PKG1 mutant mice

The above findings support the conclusion that reduced myocardial cGMP in the Db/Db and HFHS mice promotes inducible VT through reduction of PKG1α signaling, and that selective disruption of PKG1α is sufficient to promote susceptibility to VT independently of other side effects of DM or metabolic syndrome, including insulin insensitivity and autonomic dysfunction.

We next investigated potential shared molecular mechanisms downstream of PKG1α underlying the predisposition to the genesis of VT in these models. We previously demonstrated that in a mouse model for DMI, myocardial hyperactivity of GSK3β contributes to a pro-arrhythmic phenotype^13^. We determined whether GSK3β activity might also be increased in ventricular extracts of HFHS and Db/Db mice, and whether inhibition of GSK3β activity had an effect on Ca^2+^ reuptake and inducible VT. GSK3β activity is inhibited by phosphorylation at serine 9. We therefore measured myocardial pSer9-GSK3β by immunoblot in ventricular extracts from the HFHS, Db/Db and LZM models. Importantly, all 3 models displayed reduced serine 9 phosphorylated/total GSK3β compared to respective controls, indicating disinhibited, or hyperactivated GSK3β. Immunoblot data summarized in **Figure 6A** demonstrate pSer9/total GSK3β of: 0.64 ± 0.08 in HFHS-fed (n=3) vs. 1.00 ± 0.07 on control diet (n=3), p<0.05; 0.73 ± 0.06 in Db/Db (n=4) vs. 1.0 ± 0.07 in Db/+ (n=4), p=0.025; 0.35 ± 0.06 in LZM (n=3) vs. 1.00 ± 0.03 in WT (n=3), p<0.001.

**Figure 6.**
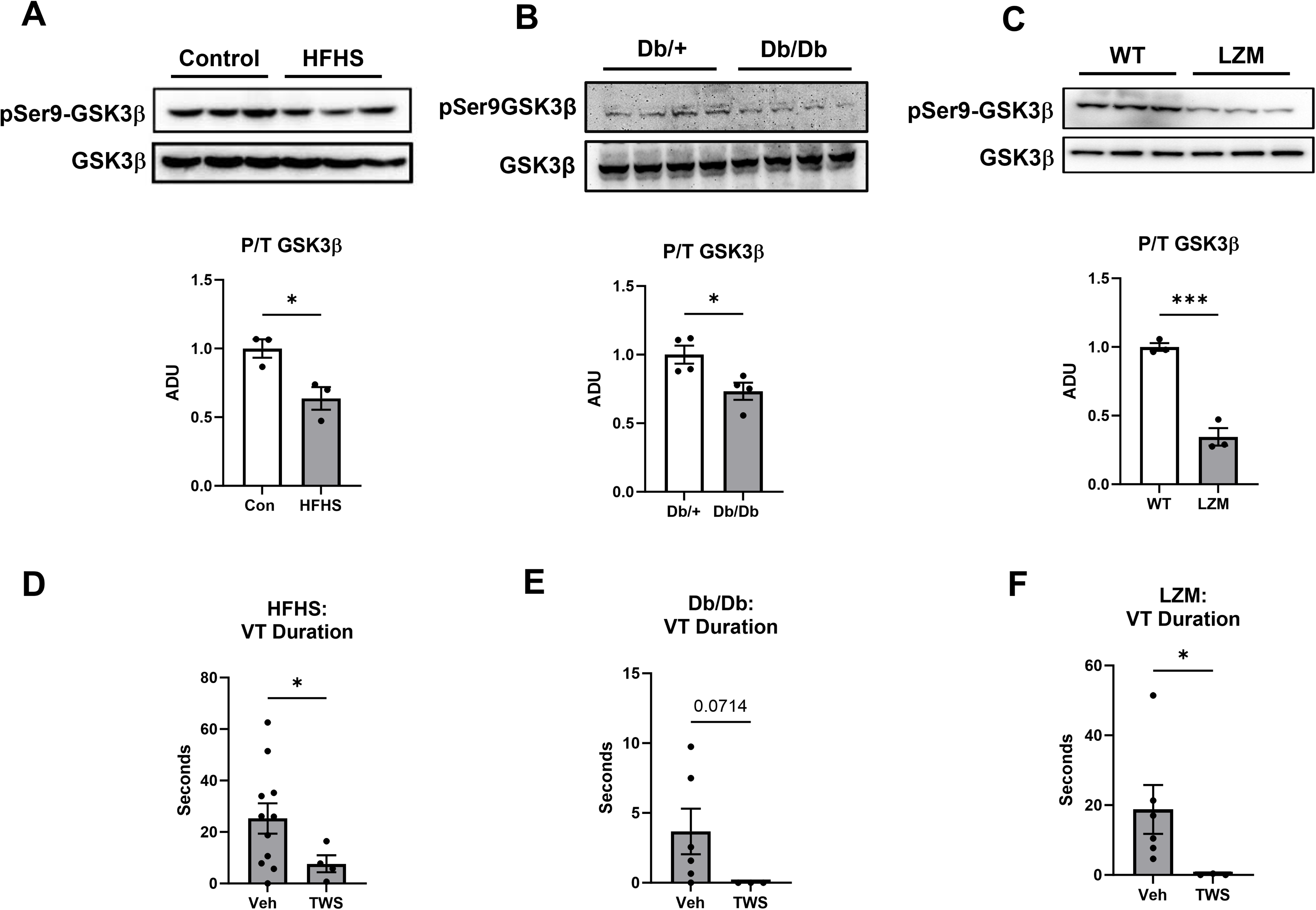
Disinhibition of GSK3β signaling in LV tissue predisposes to ventricular tachycardia in mice fed a high fat high sucrose diet, or LZM mice harboring a disrupting mutation of protein kinase G 1 alpha. Representative immunoblot and quantitative densitometry of phospho-serine 9 GSK3β normalized to total GSK3β in LV lysates from: (A) high fat high sucrose (HFHS)-fed mice; (B) Db/Db mice; and (C) PKG1α leucine zipper mutant (LZM) mice, compared with mice on control diet, Db/+, and wild type littermate controls respectively. Duration of ventricular tachycardia in response to programmed ventricular stimulation in (D) HFHS, (E) Db/Db, and (F) LZM mice treated for 7 days with either vehicle or with 20 mg/kg of the GSK3β inhibitor TWS119. *, p<0.05 and ****, p<0.001 by Student’s unpaired T test.

We next examined the effects of GSK3β inhibition on CM Ca^2+^ reuptake. Pre-treatment of HFHS-fed mice with the GSK3β inhibitor TWS119 (20 mg/kg daily via intraperitoneal injection for 7 days) decreased tau values for Ca^2+^ reuptake compared with vehicle-treated HFHS-fed mice (tau = 0.23 ± 0.02 ms, n=8, in vehicle-treated HFHS mice vs 0.17 ± 0.01 ms in HFHS-TWS119, n=7, p<0.05). Similarly, TWS099 treatment reduced tau in LZM mice compared with LZM vehicle-treated controls (tau =0.30 ± 0.03 ms in LZM vehicle-treated, n=8, vs 0.21 ± 0.02 ms in LZM-TWS119-treated mice, n=4, p<0.05). Tau values in TWS119-treated mice were not significantly different from tau values observed in cells from vehicle treated control mice (**Figure 5F**).

### Inhibition of GSK3b reduces duration of inducible VT in HFHS, Db/Db, and LZM mice

To test for a causal role of GSK3β and its associated effects on Ca^2+^ reuptake kinetics in mediating the proarrhythmogenic effects in these models, we next treated HSHS, Db/Db, and LZM mice, and their respective controls, with the selective GSK3β inhibitor TWS119. As shown in **Figure 6B**, TWS199 treatment normalized the duration of VT in response to programmed stimulation in: HFHS-fed mice (25.3 ± 5.9 sec in vehicle-treated HFHS, n=10, vs. 7.7 ± 3.3 sec in TWS119-treated HFHS, n=4, p<0.05); Db/Db mice (3.7 ± 1.6 sec, in vehicle-treated Db/Db mice, n=6, vs. 0 ± 0, n=3 in TWS119-treated Db/Db mice, p=0.07). We observed a similar reduction in LZM mice treated with TWS119, compared with vehicle-treated control LZM mice also presented in Figure 5B (18.8 ± 7.0 sec in vehicle-treated LZM mice, n=6, vs. 0.1 ± 0.1 sec in TWS119-treated LZM mice, n=3, p<0.05).

### Role of cGMP and GSK3β in abnormalities of cardiac myocyte calcium handling proteins in Db/Db, HFHS, and LZM mice

Based on the above findings of myocardial and CM calcium dyshomeostasis in the HFHS and LZM mice, we examined the expression and phosphorylation state of cardiac myocyte proteins involved in sarcoplasmic reticulum (SR) Ca^2+^ reuptake from the cytoplasm. The dephosphorylated form of the SR protein phospholamban (PLB) directly inhibits the activity of the Ca^2+^ reuptake protein sarcoplasmic/endoplasmic reticulum Ca^2+^ ATPase 2a (SERCA2a), thus impairing SR reuptake of Ca^2+^ during diastole^35^. The phosphorylation state of PLB is regulated by complex dynamics of kinases and phosphatases. PKA and by PKG1 increase inhibitory phosphorylation of PLB, leading to increased SERCA2a activity. Conversely, protein phosphatase-1 (PP1) dephosphorylates PLB resulting in increased SERCA2a activity inhibition. However, PP-1 activity is inhibited upon complexing with the inhibitor I-2^36^. GSK3β phosphorylation of I-2 on Thr72 inhibits I-2 binding to PP-1^37^. We hypothesized that the hyperactivated GSK3β in HFHS, Db/Db, and LZM myocardium would ultimately promote reduced PLB phosphorylation on Ser16/Thr17 (as outlined in **Fig. 8**). Western blot analysis of LV lysates from HFHS-fed mice, using an anti-pSer16/Thr17 PLB antibody, demonstrated markedly reduced PLB phosphorylation normalized to total PLB (pPLB/TPLB) from 1.0 ± 0.1 in extracts of control diet-fed (n=3) mice to 0.18 ± 0.03 (n=3) in extracts of vehicle treated HFHS-fed mice (p<0.05). We observed no changes in LV expression of SERCA2a or in total PLB in LV extracts of HFHS fed mice (**Figure 7A, B**). However, LV tissue from HFHS-fed mice displayed increased phosphorylation of the PP-1 inhibitor I-2 at Thr 72 (ratio of p-I2/T I-2 1.0 ± 0.2 in control diet vs. 1.6 ± 0.02 in extracts of HFHS-fed mice, p<0.05). Given that phosphorylation of I-2 at Thr72 interferes with its binding to and inhibition of PP-1, these findings support that the HFHS diet ultimately results in inhibition of SERCA2a at least in part via phosphorylation of I-2, subsequent PLB dephosphorylation via PP-1, and a resulting increase in direct PLB inhibition of SERCA2a. Importantly, phosphorylation of PLB in extracts of LV tissue of LZM mice was also decreased compared with WT controls. The ratio of Ser16/Thr17 PLB to total PLB (pPLB/TPLB) in LV lysates of LZM mice was decreased from 1.00 ± 0.02 in WT mice, to 0.41 ± 0.02 in LZM mice, (p<0.05, **Figure 7C, D**). These data support a common abnormality of the regulation of sarcoplasmic reticulum Ca^2+^ mobilization proteins in LZM and HFHS models in the pathogenesis of VT.

**Figure 7.**
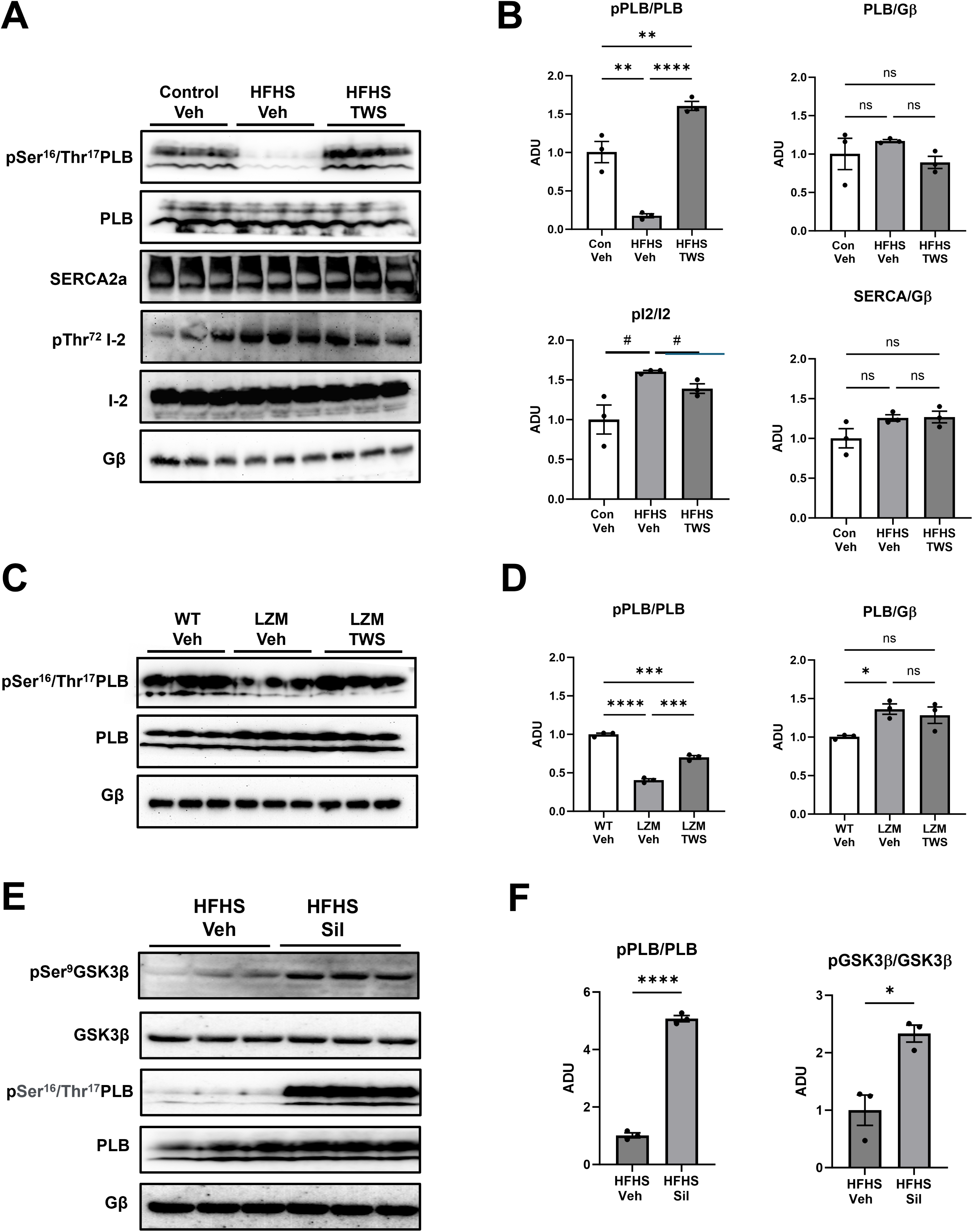
GSK3β inhibition or cGMP-pathway augmentation rescues Ca^2+^ handling protein abnormalities in LV tissue of high fat high sucrose diet and PKG1α leucine zipper mutant mice. (A) Representative immunoblots for ser16/thr17phosphorylated PLB (pPLB), phospholamban (PLB), SERCA2a, phospho-Thr72 protein phosphatase 1 inhibitor subunit 2 (pThr72I-2), I-2 and loading control Gβ in LV lysates from control or high fat high sucrose (HFHS)-fed mice, treated 7 days with vehicle or 20 mg/kg daily injection of GSK3β inhibitor TWS119. (B) Densitometry of pPLB normalized to PLB, pI2 normalized to I2, PLB and SERCA normalized to Gβ. (C) Representative immunoblot for phosphor-ser16/thr17 PLB, PLB, and Gβ in LV tissue from control or PKG1α leucine zipper mutants (LZM), treated with vehicle or 20 mg/kg daily injection the GSK3β inhibitor TWS119 for 7 days. (D) Quantitative densitometry of pPLB and PLB normalized to PLB and Gβ respectively. (E) Representative immunoblot for phospho-serine 9 GSK3β, total GSK3β, phosphor-ser16/thr17 PLB, and PLB. (F) Quantitative densitometry of phospho-serine 9 GSK3β normalized to total GSK3β and pPLB normalized to PLB. *, p<0.05; **, p<0.01; ***, p<0.001, and ****, p<0.0001 by 1-way ANOVA with Tukey’s multiple comparison tests, or Student’s unpaired T test for two group comparisons. #, P<0.05 by Student’s T test. N=3 per group.

**Figure 8.**
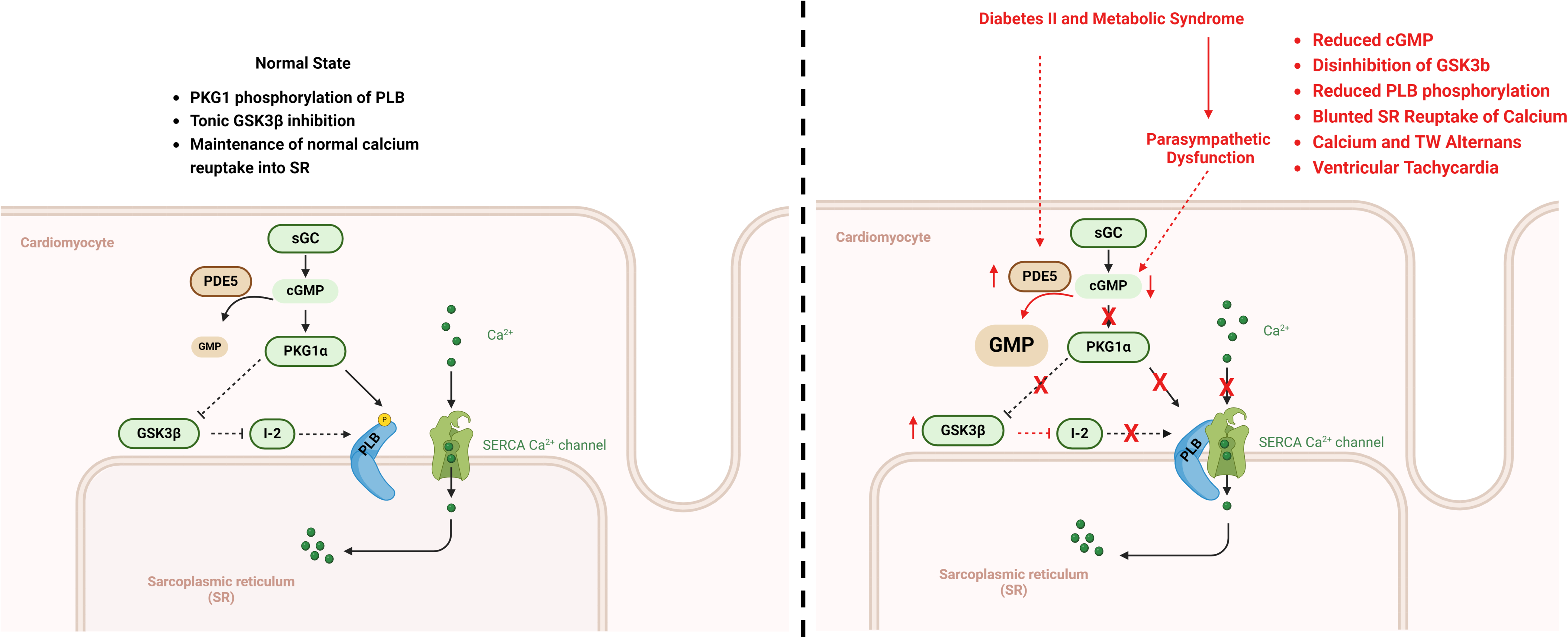
Model for the mechanism of predisposition to ventricular tachycardia in mouse models for type II diabetes and metabolic syndrome. (A) **Normal**: sGC production of cGMP activates myocardial cGMP-dependent protein kinase 1 (PKG1), leading to tonic GSK3β inhibition and preventing GSK3β-mediated inhibition of protein phosphatase inhibitor I-2. I-2 normally prevents phospholamban dephosphorylation on serine 16/17 by protein phosphatase-1 (PP-1). PKG1α directly phosphorylates phospholamban on serine 16. These dual mechanisms remove tonic phospholamban inhibition of the SERCA transporter, promoting reuptake of cytosolic calcium into the sarcoplasmic reticulum. (B) **Diabetes and metabolic syndrome:** increased PDE5 catalysis and parasympathetic dysfunction reduce available cGMP. Reduction of intracellular cGMP removes the PKG1α of GSK3β, resulting in hyperactive/disinhibited GSK3β and reduced PKG1α phospholamban phosphorylation on serine 16, thus promoting phospholamban inhibition of SERCA. The resulting blunted Ca^2+^ reuptake into the sarcoplasmic reticulum leads to EADs, Ca^2+^ alternans, T-wave alternans, and predisposition to ventricular tachycardia.

To determine directly if GSK3β kinase function played a role in these abnormalities of Ca^+2^ handling proteins, we determined the effect of the GSK3β inhibitor TWS119 on the phosphorylation of PLB. Treatment of HFHS fed mice for 7 days with TWS119 increased P/T PLB to 1.61 ± 0.05 vs. 0.18 ± 0.03 in vehicle-treated HFHS-fed mice, p<0.0001, **Figure 7A, B**.

Furthermore, TWS119 treatment of LZM mice increased P/T PLB ratio of 0.70 ± 0.02 compared with 0.41 ± 0.02 in vehicle-treated LZM mice, p<0.001 (**Figure 7B**). However, the effect of TWS119 on phosphorylation of PLB was less marked in the LZM mice than in the HFHS-fed mice. Importantly, treatment of HFHS fed mice with TWS119 had no effect on the level of SERCA2a, PLB or I-2. However, TWS119 did decrease the level of pSer72I-2/TotalI2 from 1.60 ± 0.02 in vehicle-treated HFHS-fed mice to 1.39 ± 0.06 in TWS119 treated mice, p=0.025 (**Figure 7A, B**). These data support the conclusion that increased GSK3β activity regulated Ca^2+^ reuptake at least in part via an I-2/PP-1/PLB pathway.

Given that increasing cGMP levels via sildenafil inhibition of PDE5 attenuated inducible VT in response to programmed ventricular stimulation, we determined directly whether these antiarrhythmic effects might be associated with increased inhibitory phosphorylation of GSK3β and PLB in HFHS fed mice. HFHS fed mice were treated for 1 week with sildenafil. Western blot analysis of ventricular extracts from these mice demonstrated that the ratio of pSer16/Thr17/total PLB increased from 1.01 ± 0.09 in vehicle-treated HFHS-fed mice to 5.07 ± 0.11 in sildenafil-treated HFHS-fed mice (n=3 per group, p<0.0001), while the ratio of pSer9/total GSK3β increased from 1.00 ± 0.26 in vehicle-treated HFHS-fed mice to 2.33 ± 0.15 in sildenafil-treated HFHS-fed mice (n=3 per group, p<0.05).

## Discussion

We investigated the predisposition to VT by programmed stimulation in the Db/Db and HFHS models of DMII and metabolic syndrome, respectively, to understand potential mechanisms by which these conditions promote sudden cardiac death. We observed: 1) Increased susceptibility of Db/Db and HFHS mice to inducible VT and T wave alternans in response to programmed stimulation; 2) autonomic dysfunction as evidenced by blunted heart rate sensitivity to parasympathetic stimulation, reduced cGMP, and increased PDE5 expression in LV tissue of Db/Db and HFHS mice; 3) reduction of VT in the Db/Db and HFHS mice after treatment with the cGMP augmenting drugs riociguat or sildenafil; 4) recapitulation of inducible VT and calcium dyshomeostasis in PKG1α LZM mice which have blunted response to cGMP signaling but do not have DM or metabolic syndrome; and 5) evidence of disinhibition of GSK3β activity in hearts of the above 3 models, as well as normalization of inducibility of VT and of calcium dyshomeostasis in response to GSK3β inhibition and cGMP augmenting drugs. Taken together, we interpret these findings to support that DMII and metabolic syndrome create an arrhythmic substrate through reduction in cGMP and thus reduced signaling via the cGMP effector PKG1α, leading to disinhibition of GSK3β and resultant Ca^2+^ dyshomeostasis, EADs and predisposition to ventricular tachycardia (**Figure 8**). These findings also represent, to our knowledge, the first direct demonstration that PKG1α opposes the pathogenesis of VT.

We observed a high burden of inducible VT in response to programmed stimulation in both the Db/Db and the HFHS models. Previous studies have identified increased ventricular arrhythmias in various models of DMII, such as the high fat diet model^11^. The Db/Db mouse harbors mutations in the leptin receptor, leading to obesity and DMII. Isolated hearts from Db/Db mice display increased ventricular arrhythmia during *ex vivo* programmed stimulation^10^. Our current findings now identify increased *in vivo* susceptibility of the Db/Db model to programmed stimulation-induced VT. This is important because programmed stimulation in human patients predicts VT and SCD^26,27^, and has been used to stratify patients with heart failure and reduced LV ejection fraction (HFrEF) for implantation of automatic internal cardiac defibrillators^38^. Our findings therefore support the Db/Db model as a clinically relevant tool for the study of mechanisms of VT genesis.

The HFHS diet produces a similar phenotype of obesity, but leads to insulin resistance and metabolic syndrome, rather than overt DMII^17^. Impaired glucose tolerance and metabolic syndrome, like DMII, increase SCD risk in patients^4–6^. Thus, the presence of VT inducibility in the HFHS mice holds similar clinical relevance as described for the Db/Db mice. Neither the Db/Db model nor the HFHS model develops coronary disease or acute MI. Thus, these models enable the investigation of mechanisms of VT predisposition independent from acute ischemia. More specifically, our findings in the Db/Db and the HFHS mouse models support the concept that even in the absence of coronary disease, DMII and metabolic syndrome serve as arrhythmic substrates which, in response to triggers, may lead to VT and SCD.

A principal finding of our study is the observation of shared reduction of myocardial cGMP in the HFHS and Db/Db models, and the demonstration that cGMP augmentation with the soluble guanylate cyclase stimulator riociguat or the PDE5 inhibitor sildenafil prevented VT inducibility. We observed evidence of myocardial parasympathetic downregulation in HFHS and Db/Db mice (**Figure 2**), consistent with established autonomic dysfunction in DM, diabetic autonomic neuropathy^4^. Because myocardial parasympathetic stimulation promotes cGMP synthesis^39^, we investigated myocardial cGMP levels in the Db/Db and HFHS mouse hearts. Several other lines of evidence also suggested a potential role of myocardial cGMP dysregulation in the pathogenesis of VT in DM. First, obesity and metabolic abnormalities reduce availability of natriuretic peptides (NPs)^40^, which normally promote cGMP synthesis through activation of membrane NP receptor guanylate cyclases. Second, others have observed reductions in myocardial cGMP in HFpEF, which itself often arises in the settings of obesity, metabolic syndrome or DM^41^. These varied observations led us to investigate whether dysregulation of cGMP in the LV of the Db/Db and HFHS mice promotes the VT substrate observed in DM and metabolic syndrome. Our finding that systemic cGMP augmentation with riociguat or sildenafil normalized the burden of inducible VT in the Db/Db and the HFHS mice supports a causal role of cGMP catabolism in producing the VT substrate in Db/Db and HFHS mice. Collectively, we interpret these findings to indicate that in DMII and metabolic syndrome, parasympathetic dysfunction and increased myocardial PDE5 expression lead to reduced availability of cGMP and subsequent predisposition to VT.

Our findings also provide direct evidence that the cGMP effector PKG1α normally opposes development of VT. Specifically, mutation of the cGMP effector protein kinase G 1α (PKG1α) directly increased inducibility of VT, similar to what we observed in the Db/Db and HFHS mice. In the cardiovascular system cGMP regulates several effectors including cGMP-sensitive phosphodiesterases and the cGMP-dependent protein kinase 1 (PKG1)^42^. PKG1α and 1β are encoded by the same PKG1 gene, but differ only in alternatively spliced N-terminal sequences^30^. cGMP binds PKG1 directly and activates its kinase function^30^. PKG1α contains an n-terminal, isoform-specific leucine zipper domain which mediates PKG1α protein-protein binding to critical substrates both in the vascular smooth muscle^43^ and in the cardiac myocyte^44^. Mice harboring genetic disruption of the PKG1α LZ domain, termed LZ mutant (LZM) mice, demonstrate reduced basal phosphorylation of established PKG1α substrates including myosin light chain phosphatase and RhoA^18^. The LZ domain also mediates PKG1α binding to the cardiac myocyte-specific PKG1α substrate cardiac myosin binding protein C^44^. We therefore used the LZM mice to investigate directly the role of PKG1α in modulating susceptibility to ventricular arrhythmia *in vivo*. Our findings of increased induced VT and Ca^2+^ alternans in the LZM mice identify that disruption of PKG1α sufficiently recapitulates the arrhythmia phenotype observed in the Db/Db and HFHS mice, independent of the complications of diabetes, glucose intolerance and autonomic dysfunction. Notably, prior work demonstrated that mutation of the PKG1α LZ domain abolished the therapeutic effects of sildenafil after transaortic constriction^32^, indicating that intact PKG1α mediated the therapeutic effects of cGMP in heart failure with reduced LV ejection fraction. Our current findings now directly demonstrate that disruption of PKG1α signaling promotes susceptibility to induced VT. Our findings now support that reduced cGMP in the Db/Db and HFHS myocardium leads to inducible VT through reduction of PKG1 activation. We conclude, therefore, that disruption of cGMP and downstream myocardial PKG1 signaling promotes the increased susceptibility of Db/Db and HFHS mice to VT. Importantly, PKG1 signaling is impaired in LV tissue from patients with DMII^45^, further supporting that cGMP-PKG1 disruption underlies predisposition to VT and SCD in humans with DM and insulin resistance.

We observed reduced serine 9 phosphorylation of GSK3β in LV lysates from Db/Db, HFHS, and LZM mice (**Figure 6**). Given that serine 9 phosphorylation inhibits GSK3β kinase function, these findings reveal disinhibition of myocardial GSK3β as a shared abnormality in the above models. In the current study, GSK3β inhibition with TWS119 normalized VT susceptibility in the Db/Db and HFHS mice. We previously demonstrated disinhibition of GSK3β as promoting VT susceptibility in the Akita mouse model of DMI^13^. However, our current findings now indicate a causal role of myocardial GSK3β disinhibition in promoting VT in DMII and metabolic syndrome. Further, NO-cGMP signaling inhibits GSK3β in several cell types^46^.

Notably, we also observed GSK3β disinhibition in the myocardium of PKG1α LZM mice, and demonstrated that GSK3β inhibition reduces inducible VT in the LZM mice. We therefore interpret these collective findings to support that reduced myocardial cGMP production in DM and metabolic syndrome and resultant disruption of PKG1 signaling contributes to VT substrate through disinhibition of GSK3β.

Each of the 3 experimental models in our study demonstrated accentuated QRS/T-wave alternans (TWA) and/or reduced heart rate threshold for TWA in response to burst pacing. T-wave alternans has been associated with life threatening ventricular arrhythmias^19^ and sudden death in the setting of MI, long QT syndrome, and heart failure^20^. Mechanistically, T-wave alternans has been attributed either to a temporal or spatial dispersion of myocyte repolarization or abnormalities of Ca^2+^ cycling^21,22^. Accordingly, ventricular myocytes from the LZM mice also displayed Ca^2+^ alternans, or alternating oscillations in the amplitude of Ca^2+^ transients in response to increased pacing rates. Ca^2+^ alternans may reflect the inability of the myocyte to cycle Ca^2+^ beat to beat in response to increasing heart rate and has been attributed to abnormalities in the function of Ca^2+^ handling proteins^24,25^. Our previous work identified excess T wave alternans in a mouse model of DMI, and demonstrated that GSK3β inhibition normalized TWA in DMI mice^47^. Studies in human patients have revealed TWA as an additive risk factor for SCD and for VT^48^. Experimental studies support that TWA arises as a consequence of cardiac myocyte calcium dyshomeostasis, specifically due to reduction in diastolic Ca^2+^ reuptake in the sarcoplasmic reticulum^25^. As T wave and Ca^2+^ alternans have been closely linked to the pathogenesis of VT and SCD in heart failure^23^, we interpret our findings to reveal Ca^2+^ dyshomeostasis as a shared mechanism in models of DMII, metabolic syndrome, and PKG1 disruption.

Prior studies have established that cGMP signaling in the cardiac myocyte promotes lusitropy and calcium reuptake^35^. Mechanistically, PKG1α directly phosphorylates phospholamban (PLB), leading to increased SR Ca^2+^ reuptake^35^. Here we observed reduced PLB phosphorylation in LV tissue of LZM mice (**Figure 7**). We also detected increased time constant of Ca^2+^ reuptake and a lower Ca^2+^ alternans threshold in CMs from LZM mice, compared with their WT littermates.

These findings support directly that PKG1α contributes to maintenance of normal SR Ca^2+^ reuptake, and that disruption of normal cGMP-PKG1 signaling promotes TWA through dysregulated PLB and resultant SR dysfunction. We also note that the GSK3β inhibitor TWS119 did not completely restore a normal pattern of PLB phosphorylation in LZM LV tissue. This finding is consistent with a mechanism in which PKG1α promotes PLB phosphorylation and Ca^2+^ homeostasis both through tonic inhibition of GSK3β signaling, but also through direct phosphorylation of PLB. Furthermore, we demonstrated that riociguat and sildenafil, which attenuate inducible VT in these mice, decrease levels of GSK3β activity and reverse the decrease in PLB phosphorylation consistent with the conclusion that their anti-antiarrhythmic effect might be mediated via the normalization of GSK3β mediated Ca^2+^ dyshomeostasis.

From a translational perspective, our findings suggest that pharmacological augmentation of cGMP-PKG1 signaling might reduce VT, and thus SCD, in patients with DM or metabolic syndrome. However, reduction of cGMP occurs in additional conditions associated with SCD, which therefore suggests a broader role of dysregulation of cGMP-PKG1 signaling in promoting VT in disease states other than DM. For example, myocardial tissue from patients with heart failure with preserved LV ejection fraction (HFpEF) demonstrates reduced cGMP levels, compared with HFrEF and with nonfailing hypertrophied LV tissue^41,49^. Notably, SCD remains the highest cause of death in HFpEF, and SCD risks increase in HFpEF patients with concomitant DM^50^. The reduction of cGMP in HFpEF may occur through blunted cGMP synthesis by NO-sensitive guanylate cyclase^51^, or through increased cGMP catalysis by phosphodiesterases 5 and 9^52^. Further, sildenafil use also correlates with reduced post-MI death in patients^53^. Finally, experimental studies implicate cGMP and PKG1 signaling in the regulation of ventricular arrhythmia in models of diseases other than DM. For example, PDE5 inhibition with sildenafil opposes VT both in model of acute infarction in dogs^54^, and in a drug-induced long QT syndrome model in sheep^55^. Genetic deletion of the genes for ANP and BNP in mice leads to reduced cGMP-PKG1 signaling and promotes both sudden death after transaortic constriction, and increased isoproterenol-induced VT^56^. Our findings of increased VT inducibility in PKG1α LZM mice now provide to our knowledge the first direct evidence for the role of PKG1 in normally opposing development of VT substrate *in vivo*. Notably, the LZM mouse does not develop diabetes or obesity. Accordingly, we propose that the established disruption of cGMP signaling in disease states may ultimately promote VT and SCD through dysregulation of myocardial PKG1 activity, localization, or substrate availability. PKG1-activating therapies including sildenafil, sacubitril/valsartan, vericiguat, and nitrates are safe and are routinely prescribed to patients. Whether these agents or other investigational PKG1 activators such as PDE9 inhibitors might reduce VT events remains unknown. Thus, our findings support further investigation of cGMP-augmenting therapies to reduce SCD in a variety of conditions.

### Study limitations

Our study has several limitations. First, we focused our studies on male mice only, since we did not observe inducible VT in female mice on HFHS diet. Importantly, female Db/Db and HFHS mice have sex-specific differences in their cardiac phenotypes^57^. Future work will explore the role of cGMP-PKG1 signaling in arrhythmogenesis in females, and the mechanisms underlying the sex effect in VT predisposition in these models. Second, we only used a single model of programmed stimulation to study VT susceptibility. We therefore cannot conclude whether dysregulation of cGMP, PKG1, or GSK3β alters arrhythmia susceptibility in other pro-arrhythmic states such as myocardial infarction or HFrEF. Future studies will address these important questions.

## Conclusions

In summary, we identified abnormalities of cGMP signaling as promoting VT, Ca^2+^ dyshomeostasis, and GSK3β disinhibition in two models of DMII and metabolic syndrome in mice. These findings support the translational relevance of further study of myocardial cGMP and PKG1 signaling as protecting against VT and SCD in patients with DM.

## Acknowledgements

None

## Conflicts of interest

The authors declare no conflicts of interest.

## Funding information

This work was supported by the National Institutes of Health 1R01HL162919 to R.M.B., and 1R01HL153433 to J.B.G. and R.M.B.. The funding sources had no input into the study design, collection and interpretation of data, in the writing of the manuscript, or in the decision to submit this paper.

## AUTHOR CONTRIBUTIONS

Conceptualization, X.C., R.M.B., and J.B.G.; Methodology; Y.Z., A.T., X.C., M.J.A., G.L.M., A.A., M.A., J.H., T.P., B.W., and P.L.; Formal Analysis, Y.Z., A.T., X.C., M.J.A., G.L.M., P.L., C.M., and J.B.G.; Writing-Original Draft Preparation, R.M.B. and J.B.G; Writing-Review & Editing, X.C., A.T., Y.Z., P.L., M.J.A., G.L.M., R.M.B., and J.B.G.

## Abbreviations

cGMP: cyclic guanosine monophosphate

CM: cardiac myocyte

GSK3β: glycogen synthase kinase-3 beta

LV: left ventricle

PKG: cGMP-dependent protein kinase

## Notes

### Competing Interest Statement

The authors have declared no competing interest.

